# *JNplots*: an R package to visualize outputs from the Johnson-Neyman technique for categorical and continuous moderators, including options for phylogenetic regressions

**DOI:** 10.1101/2023.05.05.539633

**Authors:** Ken S. Toyama

## Abstract

The analysis of two-way interactions in linear models is common in the fields of ecology and evolution, being often present in allometric, macroevolutionary, and experimental studies, among others. However, the interpretation of significant interactions can be incomplete when limited to the examination of model coefficients and significance tests. The Johnson-Neyman technique represents a step forward in the interpretation of significant two-way interactions, allowing the user to examine how changes in the moderator variable, it being categorical or continuous, affect the significance of the relationship between the dependent variable and the predictor. Despite its implementation in several software since its initial development, the available options to perform the method lack certain functionality aspects, including the visualization of regions of non-significance when the moderator is categorical, the implementation of phylogenetic corrections, and more intuitive graphical outputs. Here I present the R package *JNplots*, which aims to fill gaps left by previous software regarding the calculation and visualization of regions of non-significance when fitting two-way interaction models. *JNplots* includes two basic functions which allow the user to investigate different types of interaction models, including cases where the moderator variable is categorical or continuous. The user can also specify whether the model to explore should be phylogenetically informed and choose a particular phylogenetic correlation structure to be used. Finally, the functions of *JNplots* produce plots that are largely customizable and allow a more intuitive interpretation of the interaction term. Here I provide a walkthrough on the use of *JNplots* using three different examples based on empirical data, each representing a different common scenario in which the package can be useful. Additionally, I present the different customization options for the graphical outputs of *JNplots*.

## INTRODUCTION

The analysis of two-way interactions in linear models (i.e., models of the form: *dependent variable ∽ predictor * moderator*) is common in the fields of ecology and evolution (Hilborn and Stearns, 1982; Dochtermann and Jenkins, 2011; Spake et al., 2023). For example, one might be interested in the interaction between sex and size when examining the ontogenetic allometry of a trait in a population or species (*do males and females of a given species show different allometries?*). Macroevolutionary studies also necessitate the analysis of interactions. For instance, one could be interested in testing whether a set of species follows Bergmann’s rule (i.e., a positive association between body size and latitude; Bergmann, 1847), but also whether this hypothetical relationship changes depending on the degree of precipitation these species experience (*do different levels of precipitation affect the relationship between latitude and size across species?*).

These two examples illustrate four different characteristics of models in ecology and evolutionary biology that may involve two-way interactions. First, the moderator (which is an independent variable that modulates the effect the predictor has on the dependent variable) can be (1) categorical (e.g., sex) or (2) continuous (e.g., precipitation). Next, an interaction analysis can be (3) phylogenetically-independent (e.g., the comparison of the ontogenetic allometry of males and females of the same species) or can be (4) phylogenetically-informed (e.g., the examination of the effect of precipitation on the evolutionary relationship between temperature and size across species). In the latter case, the influence of shared evolutionary history needs to be accounted for through the modification of the variance-covariance matrix of the taxa involved in the analysis (Revell, 2010; Symonds and Blomberg, 2014).

Regardless, a significant interaction term in either type of model provides evidence of an effect of the moderator on the relationship between the dependent variable and the predictor. Once this is confirmed, more information about the nature of the interaction can be obtained by looking at the model coefficients. Let us consider the first example (trait ∽ size * sex). If the interaction term is significant and presents a positive coefficient a researcher would now know that the slope of the trait ∽ size relationship is significantly higher for one of the sexes. Similarly, if we obtain a significant interaction term in the second example (size ∽ latitude * precipitation) with a negative coefficient, one could infer that the slope of the size ∽ latitude relationship becomes more negative (or less positive) as precipitation increases.

However, even with this information the significant effect of an interaction might have relevant biological implications that are not immediately obvious. For example, males might have a steeper allometric slope than females for a given trait based on our inference of a significant interaction, but this does not eliminate the possibility that males and females might not be different in shape at large (Figure 1A) or small sizes (Figure 1B), or even that they might not significantly differ in shape at any size value that is biologically relevant (Figure 1C). Moreover, in Figure 1A and 1B we cannot statistically conclude that, overall, one sex has relatively larger trait values than the other, even though the visualization of the data suggests this is the case at least for most of the size range. The reason is that the assumption of homogeneity of slopes, necessary to compare groups when performing analyses of covariance (i.e., ANCOVA), is not met when the interaction term is significant (Sokal and Rohlf, 2012). Similarly, precipitation might have a significant and negative modulatory effect on the relationship between size and latitude. However, with this information alone one cannot know the precipitation values for which the size ∽ latitude relationship is significant (Figure 1D, E). It is also possible that, despite precipitation affecting the slope of this association, the size ∽ latitude relationship stays significant for all realistic precipitation values (Figure 1F). In either case, the examination of coefficients when obtaining a significant interaction term might be insufficient to interpret an interaction model and obtain useful statistical conclusions, evidencing the need for better analytical and visualization techniques.

**Figure 1.**
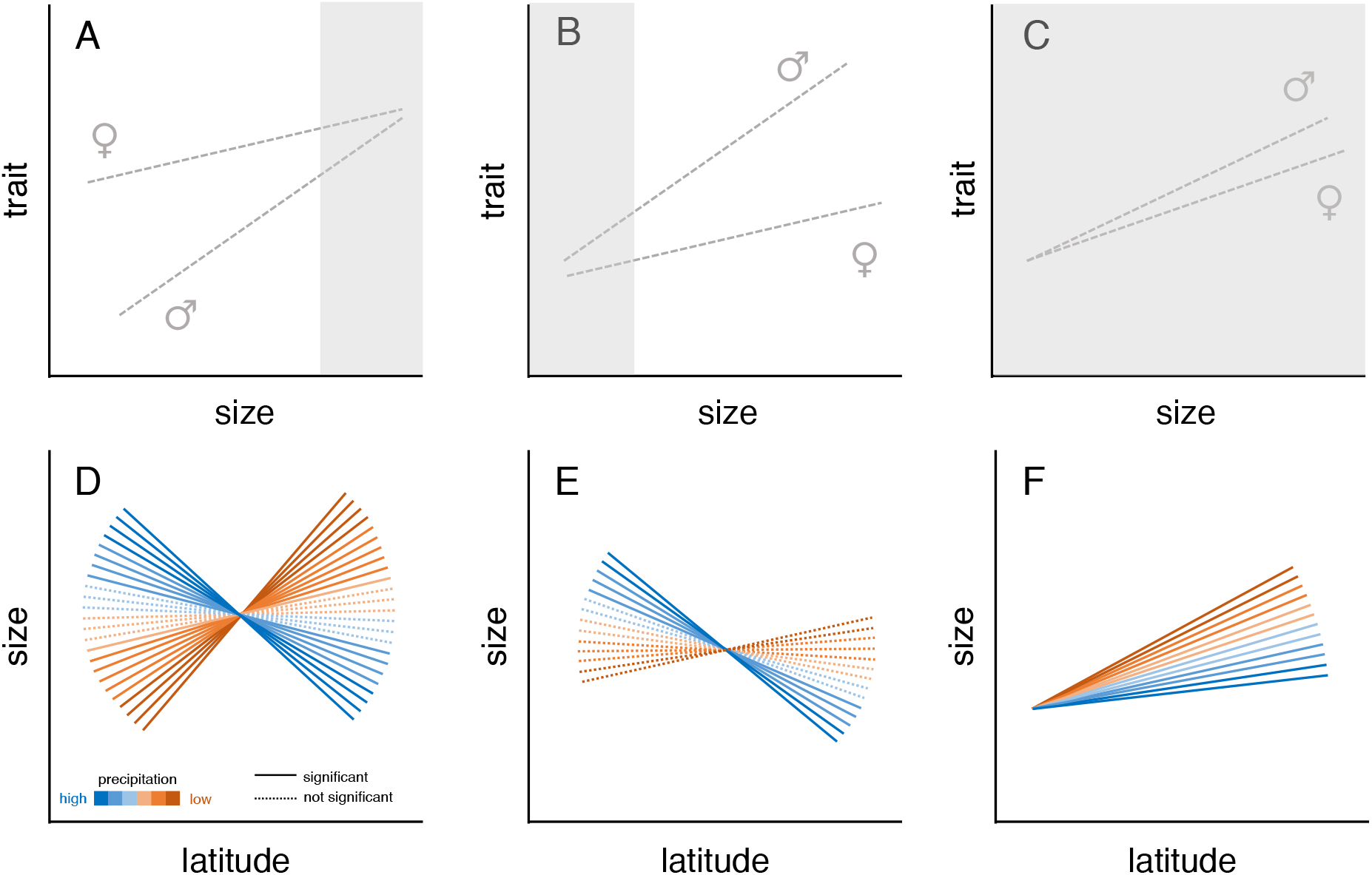
Hypothetical examples where the interactions between the moderator and the predictor are significant. A, B, and C depict cases where sex, a categorical moderator, influences the relationship between a given trait and size. In A and B, the trait value is larger in one sex than in the other, but because of the slope difference between sexes the difference in trait values might be non-significant at some unknown values of size (here depicted as grey regions). In C, the interaction between sex and size is also significant, but the different slopes do not result in significant differences between sexes for any biologically relevant value of size. In D, E, and F, precipitation, a continuous moderator, has a significant effect on the relationship between size and latitude. In D, the relationship between size and latitude is significant at high (blue) and low (brown), but not intermediate levels of precipitation (dotted lines). In E, the association between size and latitude is only significant and negative when precipitation is high. In F, the relationship between size and latitude stays positive and significant regardless of precipitation level, although it significantly influences the slope of the relationship. In D–F, solid and dotted lines represent significant and not significant size ∽ latitude associations, respectively.

### The Johnson-Neyman technique: available software, and limitations

The Johnson-Neyman technique (Johnson and Neyman, 1936; Johnson and Fay, 1950) is a method that allows a more thorough examination of interaction effects. Originally developed to account for the effect of a categorical moderator, it allows the identification of a range of predictor values for which the interaction between predictor and moderator results in non-significant differences in the dependent variable between categories (White, 2003; Huitema, 2011). For instance, it would allow one to identify size values for which trait differences between males and females are not significant in figures 1A and 1B (indicated by grey areas). The Johnson-Neyman technique has already been employed in empirical research in ecology and evolution. For example, Hünicken et al (2022) utilized the method to identify regions of non-significance in their allometric analysis of two species of *Corbicula* clams. They found the clam species showed different height ∽ length relationships (i.e., a significant *length * species* interaction). However, despite this significant interaction, the Johnson-Neyman technique indicated that the two species differed in height only at the extremes of the length distribution, while differences in height were not significant for most length values (see Figure 4D in Hünicken et al., 2022).

The Johnson-Neyman technique has also been expanded to account for continuous moderators (Bauer and Curran, 2005). Unlike the case with categorical moderators like ‘sex’ or ‘species’, one might be more interested in assessing how a gradual change in variables like precipitation or temperature affect the relationship between the dependent variable and the predictor (Figure 1D– F). For example, Jaime et al (2022) estimated the rates at which trees in 130 experimental plots were attacked by bark beetles and how these rates were affected by the climatic distance between a given plot and the niche optima of the host tree (distance_host_) and that of the beetle species (distance_beetle_) (i.e., attack rate ∽ distance_host_ * distance_beetle_). The Johnson-Neyman technique allowed the authors to conclude that, although attack rates decrease with distance_host_, this relationship weakens and even disappears as distance_beetle_ values increase (see Figure S5 in Jaime et al., 2022).

Despite being a relatively unknown method, a number of software have been developed to perform the Johnson-Neyman technique and its expanded application for continuous moderators (Preacher et al., 2006; Hayes and Matthes, 2009; Carden et al., 2017; Hayes and Montoya, 2017; Montoya, 2019; Lin, 2020), the most complete being the R (R Core Team, 2021) package *interactions* (Long, 2019), which includes all the functionality provided by other software and overall includes a wide range of visualization and analysis options. Nonetheless, this and previous software lack some functions that might prove useful for users exploring model interactions. Regarding the issue of phylogenetic relatedness, previous methods do not provide an option to directly incorporate phylogenetic information in the calculation of regions of non-significance, limiting the use of the technique in macroevolutionary studies. Regarding categorical moderators, other software do not provide an option to visualize regions of non-significance (i.e., values of the predictor for which there are no significant differences between categories, e.g., Figure 1A–C). Indeed, the uses of the Johnson-Neyman technique for categorical moderators reported in the literature are usually based on custom-made programming scripts (e.g., the study on *Corbicula* clams described above, Hünicken et al, 2022). Finally, regarding the effect of continuous moderators, the function *johnson_neyman* of the R package *interactions* provides a numerical output as well as a plot showing the association between the value of the moderator and the slope of the relationship between the dependent variable and the predictor. Although this type of plot (Figure 2A) resembles the one originally presented by Bauer and Curran (2005) and has been used to describe interaction effects in the literature (e.g., the study on bark beetles described above, Jaime et al., 2022), its interpretation is not straightforward because the relationship between the dependent variable and the predictor (e.g., as in Figure 2B) is not presented other than through the value of its slope.

**Figure 2.**
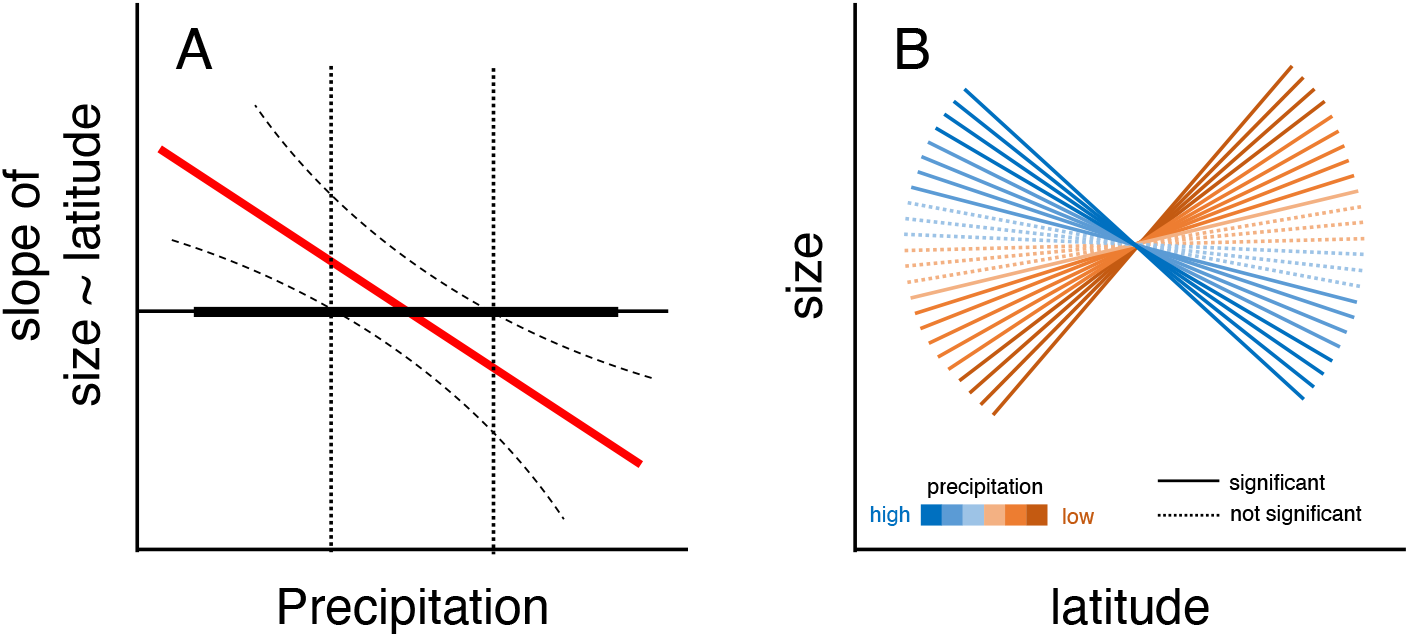
Hypothetical example of a significant interaction effect between latitude and precipitation and its effect on body size. Values of precipitation (moderator) that result in non-significant relationships between size (dependent variable) and latitude (predictor) can be obtained through an extension of the Johnson-Neyman technique. (A) The output of the method can be visualized as a plot showing the relationship between the slope of the size ∽ latitude association and precipitation values (see Bauer and Curran, 2005 and the R package *interactions*, Long, 2019). Here the horizontal thin line represents a slope of zero, the horizontal thick line represents the range of precipitation data, the red line shows the negative relationship between the size ∽ latitude slope and precipitation, the dashed lines represent 95% confidence intervals, and the vertical dotted lines represent the range of precipitation values that result in a non-significant size ∽ latitude relationship. (B) Alternatively, the different slopes of this relationship could be illustrated in a size ∽ latitude plot, showing how the relationship between size and latitude changes under the effect of different precipitation values. Both types of plots show the same information (higher precipitation decreases the value of the size ∽ latitude slope, but intermediate values of precipitation result in non-significant relationships), but B is easier to interpret.

Here I present the R package *JNplots* (Toyama, 2023) as a solution to fill gaps left by previous software regarding the calculation and visualization of non-significance regions through the Johnson-Neyman technique. As will be explained next, *JNplots* allows the user to calculate Johnson-Neyman intervals when including categorical or continuous moderators in interaction models, and to produce graphical outputs that depict them in an intuitive way. It also allows the user to modify the correlation structure of the data, allowing the consideration of phylogenetic relationships when calculating Johnson-Neyman intervals.

### JNplots: IMPLEMENTATION AND EXAMPLES

The *JNplots* R package can be used to analyse two-way interaction models that exhibit any of the four characteristics presented above (and their combinations) using the Johnson-Neyman technique and its variants. Its two basic functions, *jnt_cat* and *jnt_cont*, can be used to explore two-way interactions in which the moderator is categorical or continuous, respectively. Both functions allow the analysis of phylogenetically-informed models through the use of the function *gls* from the package *nlme* (Pinheiro et al., 2017). Both functions can be used to calculate and visualize ‘regions of significance’ in an intuitive way. Finally, the functions allow plotting flexibility as they include arguments that are passed on to the R base plot function. *JNplots* is freely available on the CRAN package repository (https://cran.r-project.org/web/packages/JNplots/) and depends on the packages *ape* (Paradis and Schliep, 2019), *nlme* (Pinheiro et al., 2017), and *scales* (Wickham and Seidel, 2022), which are downloaded from CRAN during the installation of *JNplots*. The package can be installed from CRAN and loaded using the following commands:

~~~
install.packages(‘JNplots’)
library(JNplots)
~~~

Using the following empirical examples, I present possible scenarios in which the functions from *JNplots* can be used and provide a detailed walkthrough of their implementation. All the data needed to reproduce these examples are publicly available from their respective sources and are also included in the installation of *JNplots*.

### Example 1: head length allometry in two lizard species

Data from this example comes from the study of Toyama et al (2018). In the original study the authors tested whether ontogenetic changes in the diet of a lizard (from insectivory to herbivory) corresponded to changes in its morphology (from slender to robust heads). As part of their analyses, the authors compared the head shape allometry of the semi-herbivorous species to other congeners that showed mainly insectivorous habits throughout their life (see Figure 4 in Toyama et al., 2018). Using their original data, I compared the relationship between head length and body size in a pair of these species: *Microlophus thoracicus*, a semi-herbivore species, and *M. peruvianus*, a species that rarely includes plant material in its diet.

I prepared a subset of the original dataset (dataset ‘microlophus’, included in *JNplots*), which included data on body size (measured as SVL (snout-vent-length) in millimetres), head length (also in mm), and species. Measurements were log-transformed. Since the moderator in this case is categorical (i.e., species), I proceeded to test for a possible two-way interaction between species and size (i.e., head length ∽ size * species) using the function *jnt_cat*. The only necessary arguments in *jnt_cat* are the names of the predictor (X), the dependent variable (Y), and the moderator (m). They are added to the function as character strings. The dataset also needs to be specified:

~~~
jnt_cat(X=‘svl’, Y=‘hl’, m=‘species’, data=microlophus)
~~~

Notice that the character strings must coincide with the column names in the dataset ‘microlophus’. These four arguments are the minimum needed for the function to work. The output of the function is a list that includes the summary table of the two-way interaction model (head length ∽ size * species), and the lower and upper limits of the region of non-significance (i.e., values of the predictor for which the difference between categories is not significant) (Table 1). The function also produces a plot showing the association between the dependent variable (e.g., head length) and the predictor (e.g., size), with the two categories (e.g., species) represented by different symbols and/or colors, and regression lines for each individual category based on the output of the interaction model (Figure 3).

**Table 1.** Two-way interaction fitted models obtained with *JNplots* for three empirical examples. Significant p-values are shown in bold. The limits of significance obtained using the Johnson-Neyman technique (*min JN value* and *max JN value*) and the minimum and maximum values found in the data (*min value data* and *max value data*) are shown at the bottom of each table. In the first example the moderator is categorical and these limits refer to values of the predictor (e.g., for which predictor values are the differences between moderator categories non-significant?), while in the second and third examples the moderator is continuous and these limits refer to values of the moderator (e.g., for which moderator values is the relationship between the dependent variable and the predictor (non)significant?).

|  | Coefficient | t-value | p-value |
| --- | --- | --- | --- |
| Intercept | -1.79 | -6.83 | **<0.001** |
| log(SVL) | 1.06 | 17.57 | **<0.001** |
| species | 1.59 | 5.47 | **<0.001** |
| log(SVL) x species | -0.38 | -5.50 | **<0.001** |
| min JN value | max JN value | min value data | max value data |
| 4.133 | 4.326 | — | — |

**Lizard home range**
|  | Coefficient | t-value | p-value |
| --- | --- | --- | --- |
| Intercept | 4.26 | 23.56 | **<0.001** |
| overlap | 1.77 | 9.11 | **<0.001** |
| network | -0.07 | -0.39 | 0.700 |
| overlap x network | 0.86 | 6.13 | **<0.001** |
| min JN value | max JN value | min value data | max value data |
| -3.296 | -1.360 | -2.169 | 2.481 |

**Bird coloration**
|  | Coefficient | t-value | p-value |
| --- | --- | --- | --- |
| Intercept | 0.32 | 18.40 | **<0.001** |
| precipitation | -5.03E-05 | -5.13 | **<0.001** |
| temperature | -4.40E-06 | -0.007 | 0.9947 |
| precip. x temp. | 1.30E-06 | 2.80 | **0.0056** |
| min JN value | max JN value | min value data | max value data |
| 31.106 | 84.853 | 1.7 | 27.5 |

**Figure 3.**
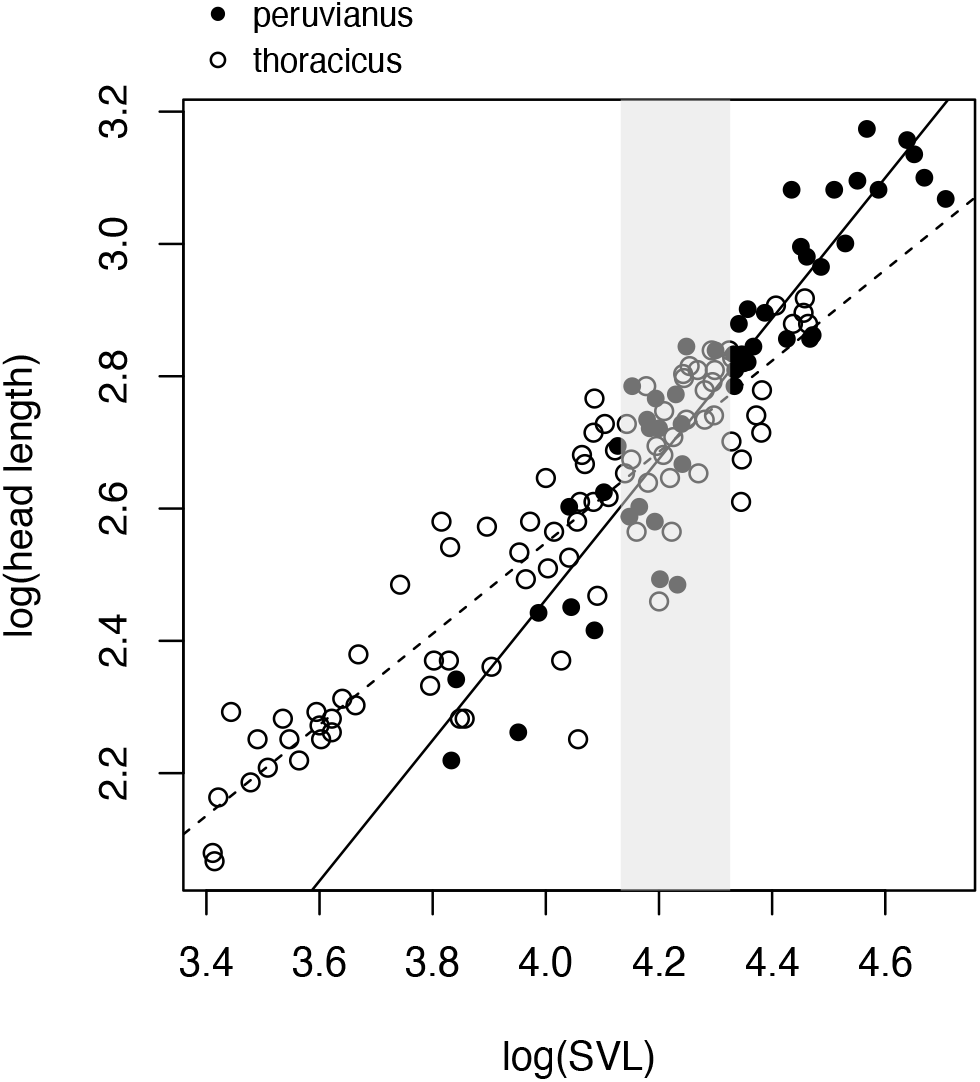
Graphical output of a model relating head length to body size (SVL) and its interaction with species of *Microlophus* lizards (model: head length ∽ body size * species) obtained with the function *jnt_cat* from *JNplots*. Solid and dashed lines represent head length ∽ body size relationships for individuals of each of the two species (also represented by closed and open circles), as shown in the legend. These relationships were obtained from the output of the interaction model. Grey area represents the non-significance area calculated with the Johnson-Neyman technique. Data obtained from Toyama et al (2018) and available to use with *JNplots*.

**Figure 4.**
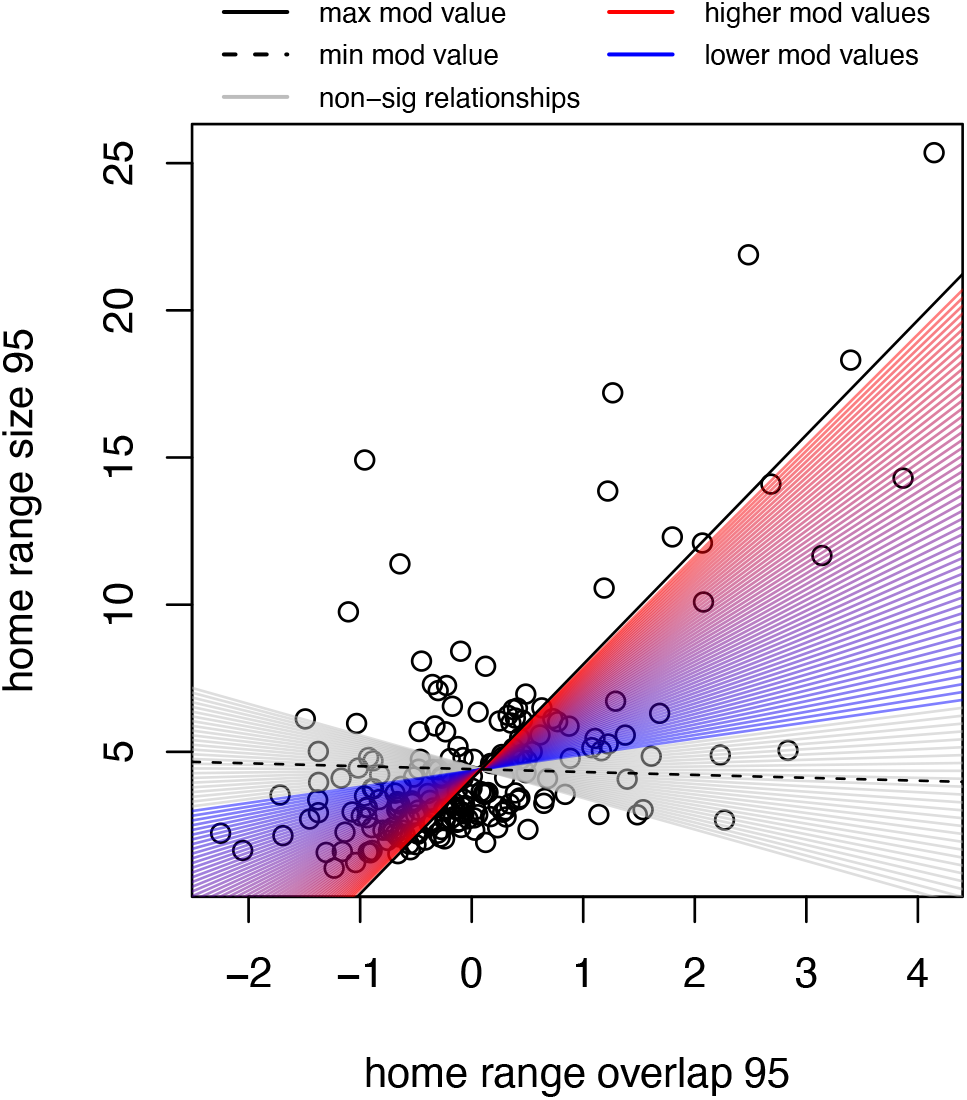
Graphical output of a model relating home range size to home range overlap and degree of social network in the lizard *Tiliqua rugosa* (model: home range size ∽ overlap * social network) obtained with the function *jnt_cont* from *JNplots*. Colored lines represent significant linear models obtained using different ‘degree of social network’ values, the blue-red gradient represents different degrees of social network going from low to high, respectively. Grey lines represent non-significant linear models. Solid and dashed black lines represent the maximum and minimum precipitation values from the dataset, respectively. Data obtained from Payne et al (2022b).

This re-analysis of the data using *jnt_cat* indicated that the interaction between sex and size was significant (*t* = -5.499, *p* < 0.001), and evidenced the existence of a region of non-significance along the examined size range (Figure 3). Specifically, the results indicated that the head lengths of both species are not significantly different for individuals with log(SVL) values between 4.13 and 4.33 (approximately between 62.33 and 75.63 mm).

In this particular case, the calculation of regions of non-significance using *jnt_cat* provided predictor (size) values that defined regions where differences between categories (and lack thereof) are statistically supported, which provides more rigor when interpreting the results of an interaction model. At this point is worth mentioning that regions of non-significance can exist and be relevant for the data of study despite the interaction term being non-significant (Rogosa, 1980, 1981; Bauer and Curran, 2005), thus it might be worth performing the Johnson-Neyman technique regardless of the significance of the interaction term.

### Example 2: drivers of home range size in a lizard

Data from this example comes from the study of Payne et al (2022a). In the original study, the authors were interested in uncovering the factors influencing the home range size (i.e., the area in which an individual interacts with the environment) of individuals of the lizard species *Tiliqua rugosa*. One of their main results indicated that the size of the home range of an individual increases with its degree of overlap with the home range of their neighbours. Additionally, this relationship is stronger for individuals that interact with more neighbours (i.e., degree of social network).

I prepared a subset of the data (dataset ‘lizard_home_range’, included in *JNplots*, see Payne et al., 2022b for original dataset) that included information on the home range size of each individual (‘hrsize95’), degree of overlap (‘PHR95_overlap-z’), and social network degree (‘degree_z’). To analyze the two-way interaction between overlap and degree of social network (i.e., home range size ∽ overlap * social network) I used the function *jnt_cont*, as the moderator (i.e., degree of social network) is continuous. As with *jnt_cat*, the necessary arguments for the function are the names of the predictor (X), the dependent variable (Y), and the moderator (m) as they appear in the dataset, which also needs to be specified:

~~~
jnt_cont(X=‘PHR95_overlap_z’, Y=‘hrsize95’, m=‘degree_z’,
     data=lizard_home_range)
~~~

As with *jnt_cat*, the output of the function is a list that includes the summary table of the two-way interaction model (home range size ∽ overlap * social network), the values of the moderator that represent the limits between significance and non-significance, and also the minimum and maximum moderator values in the data (Table 1). The function also produces a plot showing the association between the dependent variable (e.g., home range size) and the predictor (e.g., overlap) (Figure 4). However, when the moderator is continuous the interpretation of the interaction effect differs from the output of *jnt_cat*. In this example, and in agreement with the original study, home range size increases with overlap. However, the degree of social interactions has a positive effect on this relationship (i.e., the positive effect of overlap on home range size is stronger for lizards that interact more with neighbours). This positive effect is represented by multiple regression lines plotted in the output figure (Figure 4). The multiple grey regression lines that constitute the grey ‘area’ represent regressions fitted using moderator values that are outside the range of significance, i.e., values of the moderator that make the relationship between the dependent variable and the predictor not significant. (Figure 4). The ‘area’ in color consists of multiple regression lines that represent models fitted using moderator values that fall within the significance range (i.e., moderator values for which the relationship between the dependent variable and the predictor is significant). The significant regression lines are colored in a blue-red gradient that represent lower and higher moderator values, respectively, illustrating how changes in the magnitude of the moderator (i.e., degree of social network) affect the relationship between home range size and overlap (Figure 4). The plot also shows two additional lines. The solid and dashed black lines represent fitted models that use the maximum and minimum values of the moderator in the data, respectively. This aids in the interpretation of the plot because not all moderator values might be relevant for the study system or the data at hand.

In this example, a higher degree of social interactions (moderator) increases the slope between home range size (dependent variable) and overlap (predictor) (Figure 4). However, a low degree of social interactions (specifically below a value of -1.360, Table 1) makes that relationship not significant, keeping home range size small regardless of the degree of overlap (grey area in Figure 4). Importantly, some moderator values that would result in non-significant relationships are found in the data, suggesting that this result is biologically relevant (see grey regression lines between solid and dashed black lines in Figure 4).

### Example 3: drivers of coloration in birds

Data from this example were originally described in a study by Marcondes and Brumfield (2019) and reanalysed in a follow-up study (Marcondes et al., 2021). In the latter study, the authors assessed how climatic variables and light environments influence the plumage coloration of bird species of the family Furnariidae. Among other findings, the authors found that the brightness (a proxy for overall melanin content, with less bright plumage patches having less melanin) of the back plumage was negatively related to precipitation. Furthermore, an interaction between temperature and precipitation was detected, indicating that the negative effect of precipitation on brightness is stronger when temperature is lower (see Figure 1A in Marcondes et al., 2021).

I reanalysed a subset of the data used by Marcondes et al (2021) (dataset ‘birds_colors’, included in *JNplots*, original data by Marcondes and Brumfield, 2021 and Seeholzer et al., 2017) using the *jnt_cont* function as in the previous example. The model of interest in this case was brightness ∽ precipitation * temperature. However, in contrast to the previous example, this analysis implies the non-independence of datapoints due to phylogenetic relationships. To account for this, I used the argument ‘correlation’ in *jnt_cont*. The argument ‘correlation’ specifies the correlation structure of the model (as one would do in the *gls* function of *nlme*). Phylogenetic correlation structures (e.g., ‘corBrownian’, ‘corPagel’, ‘corBlomberg’, etc) in turn need a phylogeny to be specified. Here I chose ‘corPagel’ as the correlation structure and used a phylogenetic tree of the Furnariidae (‘tree_Furnariidae’, also included in *JNplots*, Harvey et al., 2020), selecting ‘1’ as the initial value of lambda (see Paradis and Schlieb, 2019, for details on using different correlation structures):

~~~
jnt_cont(X=‘bio12’, Y=‘back_bright’, m=‘bio1’, data=bird_colors,
       correlation=corPagel(1, tree_Furnariidae))
~~~

The output of *jnt_cont* showed that, in agreement with the original study, plumage brightness decreased with precipitation and the interaction between temperature and precipitation was significant (Table 1). Specifically, the effect of precipitation on brightness was stronger at lower temperatures. The limits of significance represented in the plot confirmed this pattern and also showed that the statement is generalizable for the entire range of temperature values experienced by species in the data, as it completely overlaps with the region of significance (Figure 5).

**Figure 5.**
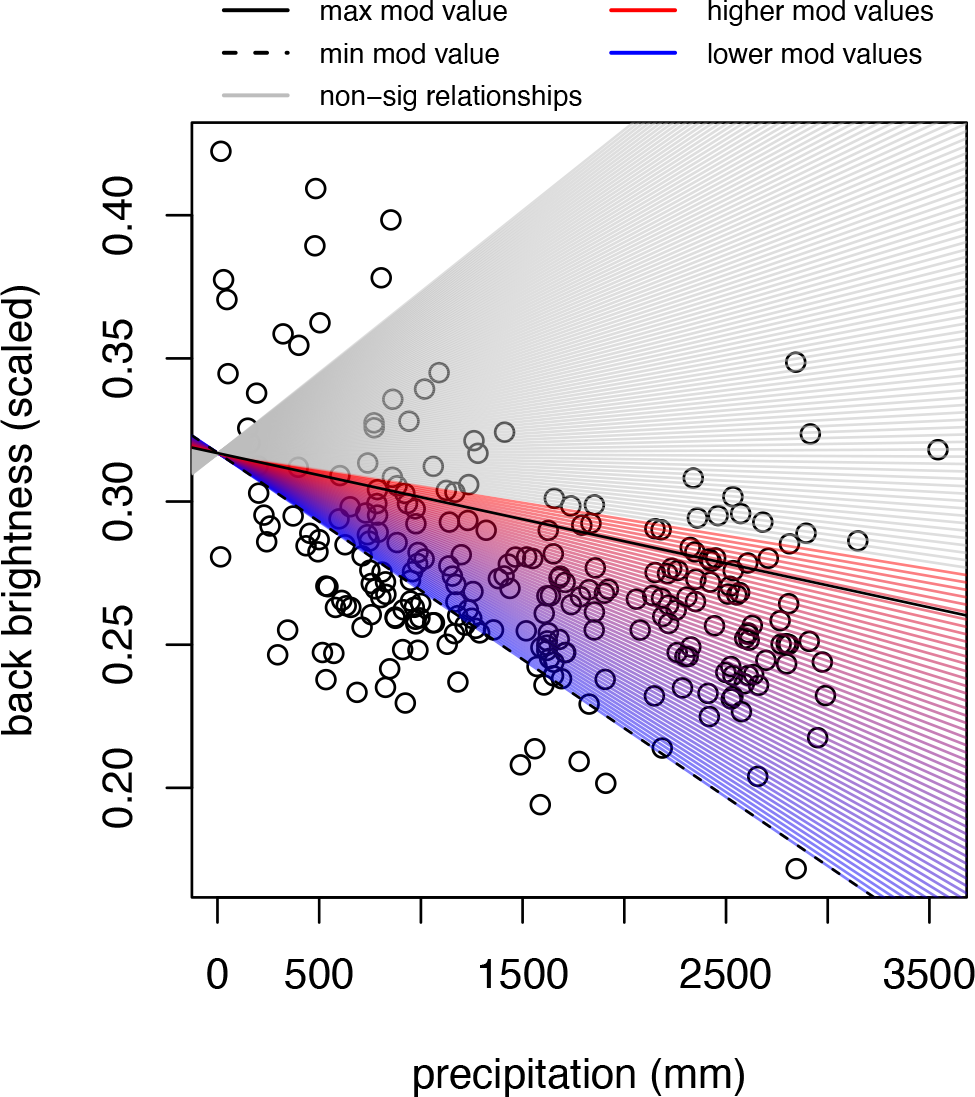
Graphical output of a model relating back plumage brightness to precipitation and temperature in Furnariidae bird species (model: brightness ∽ precipitation * temperature) obtained with the function *jnt_cont* from *JNplots*. Colored lines represent significant linear models obtained using different temperature values, the blue-red gradient represents different temperatures going from low to high, respectively. Grey lines represent non-significant linear models. Solid and dashed black lines represent the maximum and minimum precipitation values from the dataset, respectively. Data obtained from Marcondes and Brumfield (2021).

## CUSTOMIZATION OF GRAPHICAL OUTPUTS IN *JNplots*

One of the main aims of *JNplots* is to provide graphical outcomes that allow the user to interpret interaction models in an intuitive way. To aid in this objective, the graphical outputs of its functions allow for some aesthetic flexibility.

In the case of *jnt_cat* the regions of non-significance might not overlap the predictor values in the data. This would result in the region of non-significance not appearing or only partially appearing in the graphical output. Specifying the option ‘plot.full = TRUE’ (‘plot.full’ defaults to FALSE) will result in the plot always showing the entire region of non-significance regardless of its overlap with the predictor values of the data (compare Figure 6A and 6B). Other basic aspects of the plot that can be modified are the symbols representing both categories (default: pch = c(16,1)), colors (default: cols = c(‘black’, ‘black’)), line types (default: lty = c(1,2)), line widths (default: lwd = c(1,1)), and line colors (default: line.col = c(‘black’, ‘black’)). As an example, compare Figure 6A, which uses only default settings, and Figure 6C.

**Figure 6.**
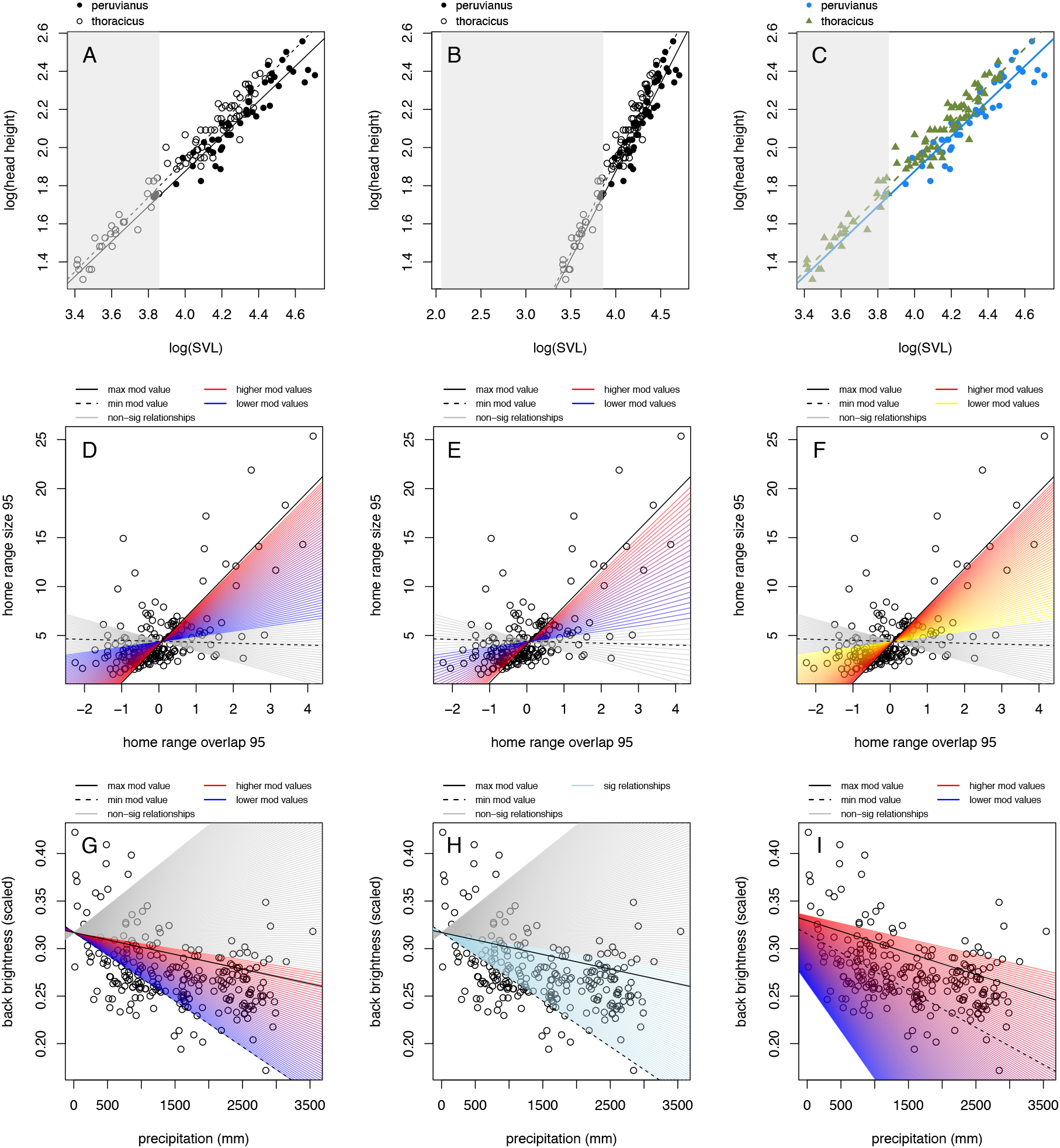
Graphical flexibility of *JNplots*. (A) *jnt_cat* was applied to a head height ∽ size * species model. (B) Here the argument ‘plot.full’ was changed to ‘TRUE’, which allows to see the entirety of the non-significance region regardless of the range of the predictor values. (C) Here plot.full = FALSE, but other arguments were modified to change the aesthetics of the plot (pch = c(16,17), cols = c(‘dodgerblue2’, ‘darkolivegreen4’), lwd = c(2,2), line.col = c(‘dodgerblue2’, ‘darkolivegreen4’)). (D) Same as figure 4, *jnt_cont* was applied to the model home range size ∽ overlap * network. (E) Here the argument ‘res’ was specified to be 40 (default = 100). Notice the lower number of regression lines and the larger space between them. (F). Here res = 80, and gradient colors are changed specifying min_col_grad = ‘yellow’ and max_col_grad = ‘red’. (G) Same as figure 5, *jnt_cont* was applied to the model brightness ∽ precipitation * temperature, res=150, correlation = corPagel(1, tree_Furnariidae). (H) Argument col.gradient = FALSE and sig_color = ‘lightblue’. The argument sig_color is only considered when col.gradient = FALSE and defines a single color to be used for all significant regression lines. The argument nonsig_color works similarly for non-significant regression lines. (I) In this case the correlation structure is based on a Brownian motion model of evolution (correlation = corBrownian(1, tree_Furnariidae).

Plotting characteristics can also be specified in *jnt_cont*. The user can control the relative number of regression lines to be plotted with the argument ‘res’, which defaults to 100. The exact number of lines to be plotted is equal to the value of ‘res’ – 1, meaning that the number of plotted regressions increases with the value specified in ‘res’ (Compare Figure 6D and 6E, which have ‘res’ values of 80 and 40, respectively). The gradient of colors shown by the significant regression lines can also be modified. The arguments ‘max_col_grad’ and ‘min_col_grad’ define the colors of the regression lines when considering the maximum and minimum moderator values that result in significant relationships, respectively. The colors of the regression lines in-between will form a gradient between the extreme colors (‘max_col_grad’ and ‘min_col_grad’ default to ‘red’ and ‘blue’, respectively). For example, compare Figure 6D and Figure 6F. If a color gradient indicating different moderator values is not desired then ‘col.gradient = FALSE’ (defaults to TRUE) should be specified. In this case, all the lines representing significant fitted regressions will take the color specified in the argument ‘sig_color’, which defaults to ‘lightblue’ (Compare Figure 6G and 6H). The color of the non-significant regression lines can also be specified in the argument ‘nonsig_color’ (defaults to ‘grey’).

Finally, as previously mentioned, the correlation structure of the data can be modified in both *jnt_cat* and *jnt_cont*. Although this is not an aesthetic specification, changing the correlation structure will most likely change the aspect of the graphical outcome of either function. For example, compare Figure 6G and 6I, which use Pagel’s lambda and Brownian motion correlation models, respectively.

## CONCLUSIONS

Multiple model testing is common in ecological and evolutionary studies, and understanding how variables included in such models interact is indispensable for their interpretation (Hilborn and Stearns, 1982; Dochtermann and Jenkins, 2011; Spake et al., 2023). Although the Johnson-Neyman technique was initially developed in the context of educational and psychological studies (Johnson and Neyman, 1936; Johnson and Fay, 1950), its application to other fields is evident (e.g., White, 2003), as was the need to expand its application beyond categorical moderators and two-way interactions (e.g., Bauer and Curran, 2005). In the same vein, *JNplots* aims to be a tool that facilitates the application of the method in ecological and evolutionary studies through the direct implementation of phylogenetic corrections and the possibility to analyze categorical and continuous moderators, thus going beyond what is possible with existing software. Equally important, *JNplots* aims to aid in the interpretation of two-way interactions through more intuitive graphical outputs.

Although its main functions are readily available, *JNplots* still has room for expansion. For example, the Johnson-Neyman technique can be applied to three-way or higher-level interactions (Pothoff, 1964; Hunka, 1995; Hunka and Leighton, 1997; Curran et al., 2004; Bauer and Curran, 2005). Other types of regressions, like type II or reduced major axis regressions, and even non-linear models also represent alternatives to traditional linear models not yet included as analytical options in this package. These and other variations in the analysis of interactions remain to be implemented in *JNplots* (or any other software). Before then, users interested in such variations are free to copy and modify the functions from *JNplots* (https://github.com/kenstoyama/JNplots) and adapt them to their needs.

Together with the release of this package, I provided a quick start guide online (https://kenstoyama.wordpress.com/2023/04/28/jnplots-quick-guide/) for users that are more familiar with the Johnson-Neyman technique and are specifically interested in the numerical and graphical outputs of *JNplots*. The same page can be used to report issues with the use of the package.

## ACKNOWLEDGEMENTS

I want to thank Miriam Ahmad-Gawel, Rafael Marcondes, and Liam Revell for their helpful comments and suggestions in earlier versions of this manuscript.

